# High-speed imaging of giant unilamellar vesicle formation in cDICE

**DOI:** 10.1101/2023.10.13.562183

**Authors:** Lori Van de Cauter, Yash K. Jawale, Daniel Tam, Lucia Baldauf, Lennard van Buren, Gijsje H. Koenderink, Marileen Dogterom, Kristina A. Ganzinger

## Abstract

Giant unilamellar vesicles (GUVs) are widely used as *in vitro* model membranes in biophysics and as cell-sized containers in synthetic biology. Despite their ubiquitous use, there is no one-size-fits-all method for their production. Numerous methods have been developed to meet the demanding requirements of reproducibility, reliability, and high yield, while simultaneously achieving robust encapsulation. Emulsion-based methods are often praised for their apparent simplicity and good yields; hence, methods like continuous droplet interface crossing encapsulation (cDICE) that make use of this principle, have gained popularity. However, the underlying physical principles governing the formation of GUVs in cDICE and related methods remain poorly understood. To this end, we have developed a high-speed microscopy setup that allows us to visualize GUV formation in real-time. Our experiments reveal a complex droplet formation process occurring at the capillary orifice, generating both larger droplets and, likely, GUV-sized satellite droplets. According to existing theoretical models, the oil-water interface should allow for crossing of all droplets, but based on our observations and theoretical modelling of the fluid dynamics within the system, we find a size-selective crossing of GUV-sized droplets only. Finally, we demonstrate that proteins in the inner solution affect GUV formation by increasing the viscosity and altering lipid adsorption kinetics. These results will not only contribute to a better understanding of GUV formation processes in cDICE, but ultimately also aid the development of more reliable and efficient methods for GUV production.

**Graphical abstract:** 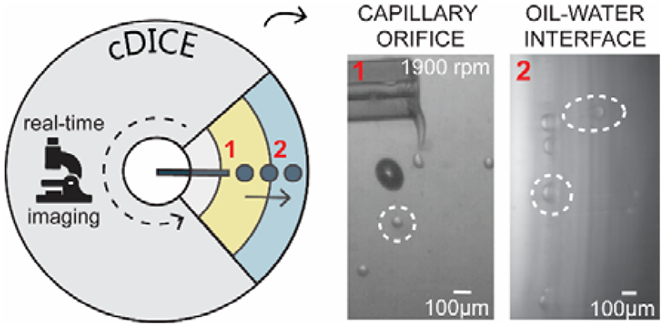

We developed a high-speed microscopy setup to study giant unilamellar vesicle formation in cDICE, revealing a complex droplet formation process occurring at the capillary orifice and size-selectivity at the oil-water interface.

## Introduction

The quest to understand and manipulate the building blocks of life, including the countless interacting molecules and biochemical reactions making up cellular life, is a major aim of biophysics and synthetic biology.^1^ One key tool in these fields are giant unilamellar vesicles (GUVs) as cell-sized, lipid bilayer-enclosed reaction compartments.^2,3^ Since their first description^4^ in 1969, GUVs have proven to be a powerful and versatile tool as they can be directly observed using real-time microscopy and easily manipulated using biophysical tools, making them ideal *in vitro* model membrane systems.^3,5,6^ More recently, GUVs have also been proposed as containers for a future synthetic cell^7–10^ and as reaction containers for chemistry and more complex cargo carriers in drug delivery^11,12^.

Despite the widespread research use of GUVs, there is still no one-size-fits-all method for their production^10^. Over the years, numerous methods have been developed to meet the demanding requirements of reproducibility, reliability, and high yield, while simultaneously achieving robust encapsulation. Historically, swelling-based methods (natural swelling^4^, electroformation^13–16^, and gel-assisted swelling^17–20^) have been used extensively for studying the biophysical properties of membranes. However, these easy-to-implement, high-yield methods offer poor control over encapsulation efficiency and the stoichiometry of encapsulated molecules. Thus, they only offer limited compatibility with establishing complex reconstituted systems. Emulsion-based techniques (water-in-oil droplets crossing an oil-water interface using gravity, centrifugation, microfluidic devices, or microfluidic jetting^21–27^) on the other hand, offer more control over GUV content and for complex encapsulation experiments. Despite the potential cost of residual membrane impurities^10,28,29^, emulsion-based methods have therefore gained popularity in recent years.

One method that particularly gained a lot of traction is called continuous droplet interface crossing encapsulate (cDICE).^30–36^ In cDICE, aqueous droplets that are produced at a capillary orifice are continuously forced through an oil-water interface by centrifugal force in a rotating chamber, thereby forming a lipid bilayer and thus GUVs.^30^ Recent optimization has made the method compatible with a wide range of biological systems, thereby offering control over encapsulated content, a high GUV yield, and straightforward implementation.^31^ However, our understanding to which degree the encapsulated contents’ complexity in cDICE can be extended, with respect to both physical properties (*e*.*g*. viscosity of encapsulated fluid) and physicochemical properties (*e*.*g*. which proteins and protein systems), remains limited. While many successes have been celebrated using cDICE, we still do not understand the underlying GUV formation process and how this affects the inherent variability in content encapsulation and yield seen in cDICE.^10^

To gain a deeper understanding of GUV formation in cDICE, we have developed a high-speed microscopy setup that allows us to visualize the GUV formation process inside the rotating chamber in real-time. We focused on the capillary orifice, where initial droplet formation occurs, and on the oil-water interface, where droplets are converted into GUVs. Our experiments reveal a complex droplet formation process occurring at the capillary orifice, governing both the formation of larger droplets and, likely, satellite droplets of the size of typical cDICE GUVs (12 µm -average diameter of GUVs formed with cDICE^31^). The transfer of these droplets through the oil-water interface appears to exhibit selectivity toward GUV-sized droplets. We support these experimental observations with scaling arguments. Finally, we demonstrate that the addition of protein to the inner solution increases the viscosity and alters the kinetics of lipid adsorption, thereby significantly influencing the process of GUV formation.

## Results and Discussion

### Design of an imaging setup to visualize droplet and GUV formation in cDICE

In the cDICE method, the initial step of GUV formation is the generation of droplets at a capillary orifice, which is inserted perpendicularly into the oil layer in the rotating chamber. In its original implementation, cDICE uses a capillary diameter of 2-25 μm to allow for tight control over GUV sizes^30^. However, we and others found such narrow capillaries to be very impractical when encapsulating protein solutions, as these capillaries are prone to rapid clogging, leading to highly irreproducible results. In our previous work, we showed that this issue can be circumvented by using wide capillaries with a diameter of 100 μm.^31^ The flow regime is therefore significantly different from the original protocol^30^, and one would not necessarily expect tight control over droplet. Still, we found that these capillaries produced a surprisingly narrow size distribution of GUV sizes, roughly ten times smaller than the capillary orifice (∼10 μm vs ∼100 μm)^31^.

To better understand how a large capillary orifice can still lead to such relatively monodispersed GUV size distribution in cDICE, we developed a high-speed microscopy setup to, for the first time, visualize the processes of droplet and GUV formation in cDICE in real-time (Fig. 1). We designed the setup so that the camera is suspended vertically above the cDICE apparatus, capturing the light of a light source located directly beneath the rotating chamber (see method section for a full description of the setup; Fig. 1a). This way, we are able to capture the process along the horizontal axis of the rotational chamber: from the capillary orifice, where initial droplet formation occurs (Fig. 1b i), to the oil-water interface, where droplets are converted into GUVs (Fig. 1b ii). Due to the high rotation speeds that are used in cDICE (∼ 1900 rpm), all processes happen on a very fast timescale, on the order of milliseconds (10^− 6^ 10^− 5^ s). To obtain a sufficiently high time resolution, we therefore used fast cameras in combination with brief exposure times up to 1 µs, reaching frame rates up to 30,000 fps.

**Figure 1.**
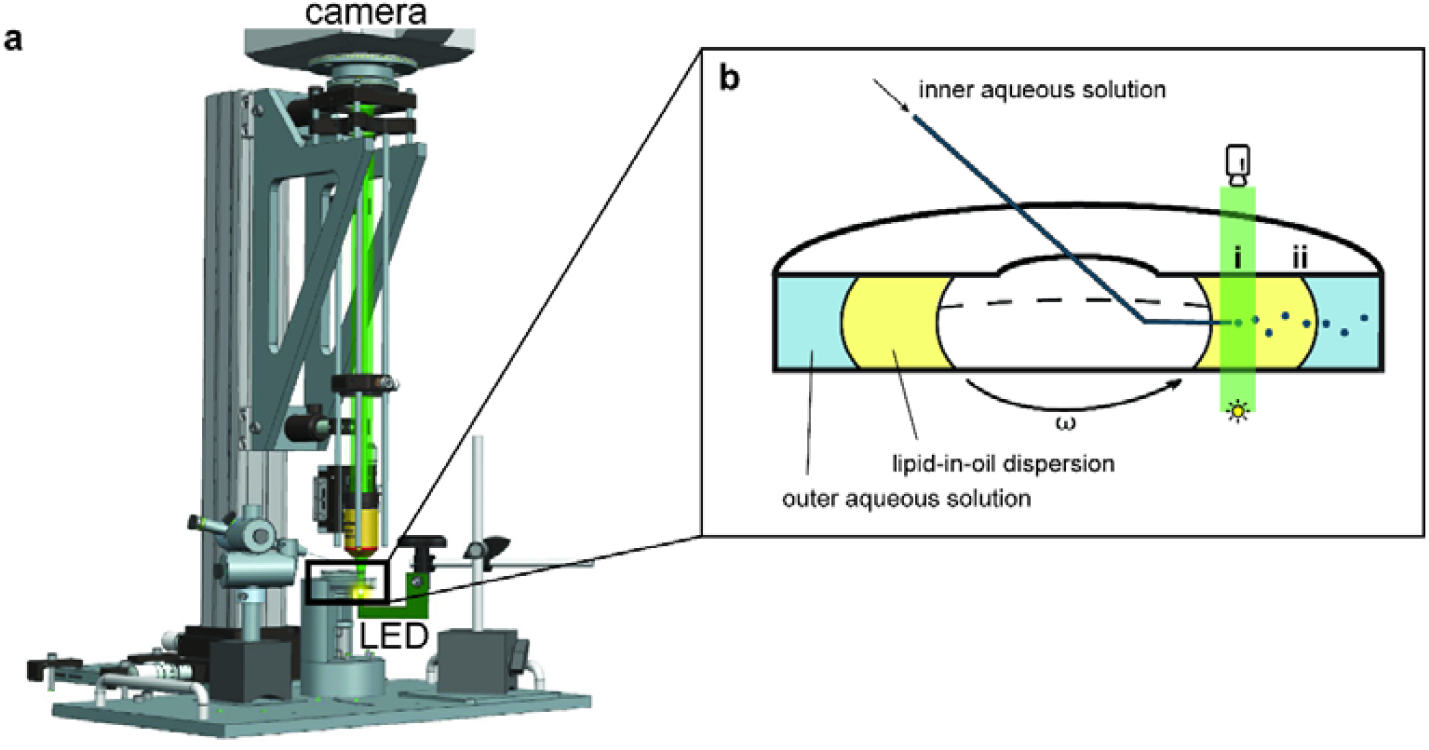
Development of a high-speed imaging setup to visualize GUV formation in cDICE. **a**. The imaging setup consists of a high-speed camera suspended above the rotating chamber and an intense light source located directly below the rotating chamber. For an interactive 360° view of the setup, see method section. **b**. In cDICE, (i) aqueous droplets are generated at the capillary orifice; subsequently travel outward through the lipid-in-oil dispersion; and finally (ii) traverse the oil-water interface, where droplets are converted into GUVs.

### Droplet formation at the capillary orifice is governed by shear forces

When we focused our imaging setup on the capillary orifice at our default conditions for GUV production (100 µm diameter fused silica capillary, a rotation speed of 1900 rpm, and a flow rate through the capillary of 25 µLmin^− 1^; see method section for further details), it immediately became clear that droplet formation under these conditions is a non-uniform, highly dynamic process with an irregular breakup pattern of a liquid filament into individual droplets (Fig. 2a, SI Mov. 1). Instead of the distinct droplet formation expected for low Reynolds numbers^30,37^, we observed fluid exiting the capillary forming a liquid filament, which often adhered to the capillary. Droplets breakup took place at the end of the liquid filament at a fast rate, with droplet sizes clearly larger than the average cDICE GUV ((68.6 ± 2.8) μm, approximately 2500 droplets per second).

**Figure 2.**
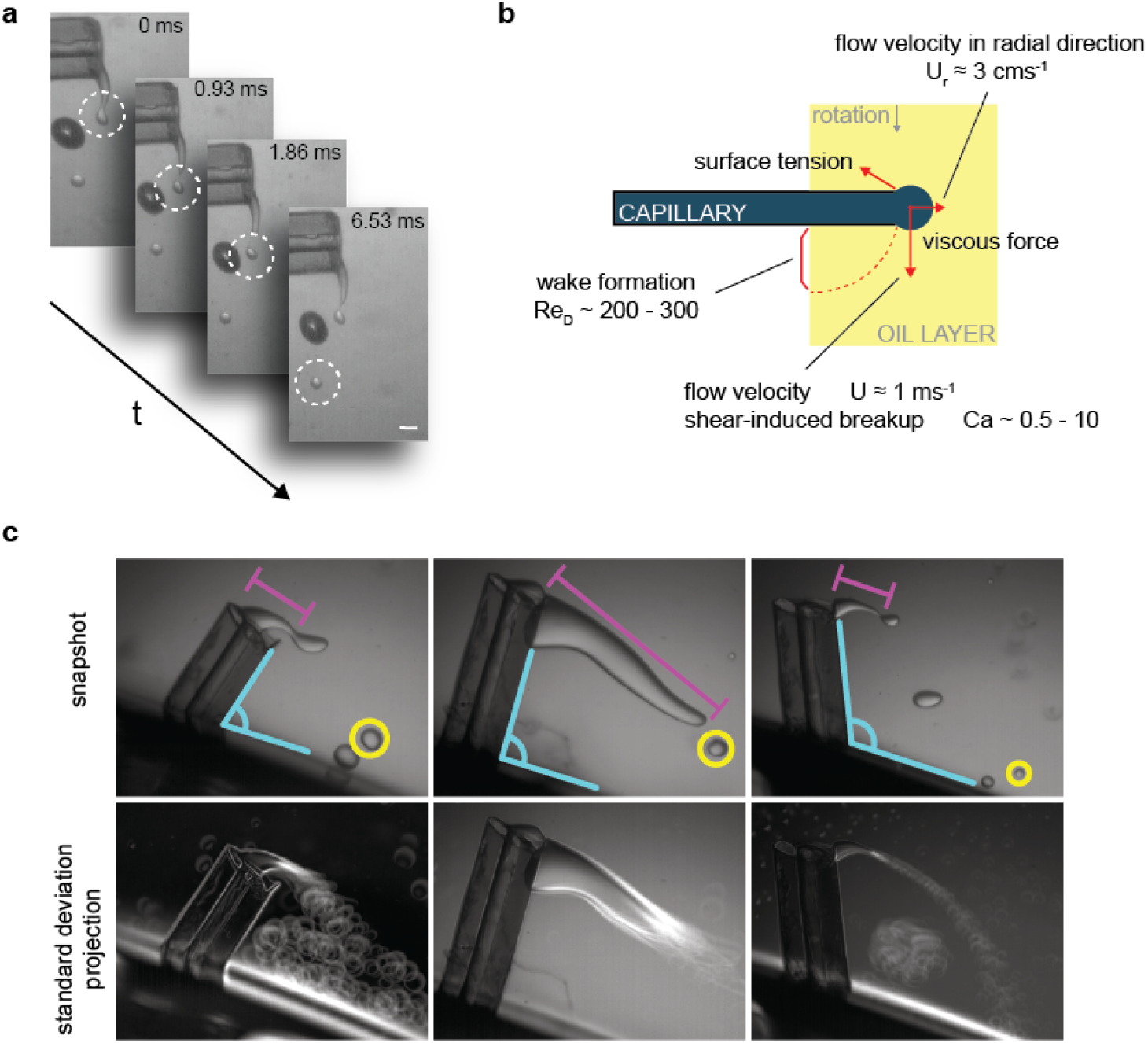
Droplet formation at the capillary orifice is governed by shear forces. **a**. Microscopic image sequence capturing a droplet of PBS buffer with 18.5 % v/v OptiPrep™ being sheared off from the liquid stream at the capillary orifice at a rotation speed of 1900 rpm. Scale bar indicates 100 μm. **b**. Illustration depicting the different forces acting at the capillary orifice: the capillary number Ca (Ca ∼ 0.5-10) indicates a shear-induced breakup mechanism, while the Reynolds number (Re_D_ ∼ 200-300) describes the wake formation behind the capillary. The shear velocity (*U* ≈ 1 ms^-1^) is larger than the flow velocity in the radial direction (U_*r*_ ≈ 3 cms^-1^), further indicating droplet formation is shear-induced. c. Microscopy images highlighting the high inter-experiment variability using same capillary in independent, consecutive experiments using the same capillary. For identical experimental conditions, noticeable differences can be seen, *e*.*g*. in insertion angle (top row, cyan), liquid filament (top row, magenta), and droplet size (top row, yellow). Bottom row: standard deviation stack projection of 100 frames (every 50^th^ frame of 5000 frames). White highlights indicate variations in movement, such as the occurrence of a droplet vortex in the wake of the capillary (left and right image) and the movements of the liquid filament end (left and middle).

Upon silanization of the capillary, we no longer observed the fluid adhering to the capillary, resulting in a more regular droplet breakup mechanism (SI Fig. 1). This can likely be explained by an increased surface hydrophobicity upon silanization when compared to the default polyimide capillary coating surrounding the capillary and the uncoated cut tip cross-section with chipped coating edges, which results in less wetting of the capillary surfaces.

We observed a significant variability in the droplet breakup dynamics at the end of the liquid filament. Factors contributing to this variability include irregularities in the capillary orifices resulting from suboptimal cutting or capillary deterioration over sustained use, differences in capillary insertion angle, and the occasionally-observed presence of an air pocket at the base of the capillary (Fig. 2a,c). Note that in all these cases, the experimental condition was indistinguishable by eye, and the differences only became apparent when visualizing droplet formation with our dedicated imaging setup.

To better understand the observed droplet breakup mechanisms, we turn to scaling arguments to rationalize our findings (Fig. 2b). Our video recordings (SI Mov. 1) suggest that droplet breakup at the tip of the capillary is not due to inertial jetting, but instead is induced by viscous shear stresses. For droplets forming from a capillary of diameter *D*, inertial jetting is expected for flow rates larger than a critical flow rate scaling with ∼π *D* ^3^γ/2ρ_*i*_)^1/2^, where ρ_*i*_ denotes the density of the inner solution and γ is the interfacial tension between the dispersed and the continuous phases^37^. In our experiments, the flow rate through the capillary is 25 µLmin^− 1^, which is significantly lower than the critical flow rate. This is consistent with our observation that droplets are indeed sheared off the capillary. Here, we must therefore consider the balance between surface tension and viscous forces characterized by the capillary number *Ca. Ca* is given by *Ca* = *µU/*γ. The flow velocity *U* at the point of insertion of the capillary is *U* = Ω*R*_*i*_, where *R*_*i*_ is the distance between the capillary orifice and the center of rotation of the chamber and Ω is the rotation speed. With *R*_*i*_ ∼ 1 *cm*, Ω ∼ 1000 − 2700 *rpm, µ* ∼ 4 − 5 10^− 3^ *kgm*^− 1^*s*^− 1^ and assuming an interfacial tension between the inner solution and the oil phase of γ ∼ 10^-3^ − 10^− 2^ *mNm*^− 1 31^, the capillary number ranges between 0.5 and 10. Monodispersed droplets form at the tip of the capillary through a dripping mechanism for low values of the capillary number^30^. Within the higher range of *Ca* reached in our experiments, droplets are therefore expected to deform and the break up mechanism to be unstable, in agreement with our observations.

cDICE experiments require high angular velocities ω (> 1000 rpm), producing flow instabilities in the wake of the static capillary inserted into the rotation chamber. Indeed, the Reynolds number, characteristic of the flow around the capillary, *Re*_*D*_ = ρ*UD*/*µ*, yields values in the range of *Re*_*D*_ ∼200 -300, with ρ ∼ 0.934 *g*/*mL* being the density of the lipid-in-oil dispersion. For *Re*_*D*_ ≥ 47, periodic vortex shedding in the wake of a cylinder is expected^38^, and for *Re*_*D*_ ≥150, further three-dimensional instabilities are predicted^38^, suggesting that the wake around the capillary will also affect droplet breakup. Indeed, we observe oscillations in droplet breakup, caused by the non-linear effects in the wake, and the inner solution adhering to the outer capillary surface. Additionally, we observed that the droplets did not immediately travel outward as expected, but rather initially exhibited an inward movement in the wake of the capillary and towards the center of the rotating chamber, before travelling outward. The larger diameter of the capillary leads to a larger capillary number Ca and to a wake instability, which both contribute to a less stable droplet breakup and larger variation in droplet size compared to previous work^30^.

### Droplet size in contrast to GUV size, is dependent on the rotation speed

To explore factors that influence droplet breakup in cDICE, we next altered the rotation speed of the rotating chamber. As the rotation speed of the chamber increases, the flow velocity at the capillary orifice also increases and the viscous forces become stronger. This leads to the droplets being more likely to break up, resulting in smaller droplets. In line with this expectation, an increase in rotation speed to 2700 rpm resulted in smaller droplets formed at a higher frequency ((28.5 ± 8.7) μm and ∼34,500 droplets per second; Fig. 3, SI Mov. 2). Decreasing the rotation speed to 1000 rpm, the lowest speed at which oil and water layers maintain a vertical interface and GUVs can be produced, had the opposite effect, i.e. larger droplets formed at a lower frequency ((273 ± 41) μm and ∼40 droplets per second; Fig. 3, SI Mov. 3). We can estimate the droplet size from a force balance between the surface tension force ∼π*D* γ and the viscous force ∼6 π*µaU*, where *D* is the outer diameter of the capillary and a is the radius of the droplet^37^. The droplet size above which break up is expected, scales with the inverse of the capillary number *a*/*D* ∼ 6 *Ca*)^− 1^, and we predict a droplet diameter of ∼100µm at 1900 rpm increasing to ∼200µm when the rotation rate is decreased to 1000 rpm, in agreement with our experimental results. Droplet formation is thus shear-induced in a broad range of rotation speeds, encompassing both lower and higher speeds than the default of 1900 rpm. Our observation that droplet size is dependent on chamber rotation speed contrasts with the size distributions for GUVs obtained using these conditions: these distributions are all indistinguishable from one another and centered around 12 μm (SI Fig. 2), thus 3-30-fold smaller in diameter than the produced droplets. Hence, at least some of the droplets formed at the capillary are not directly converted into GUVs.

**Figure 3.**
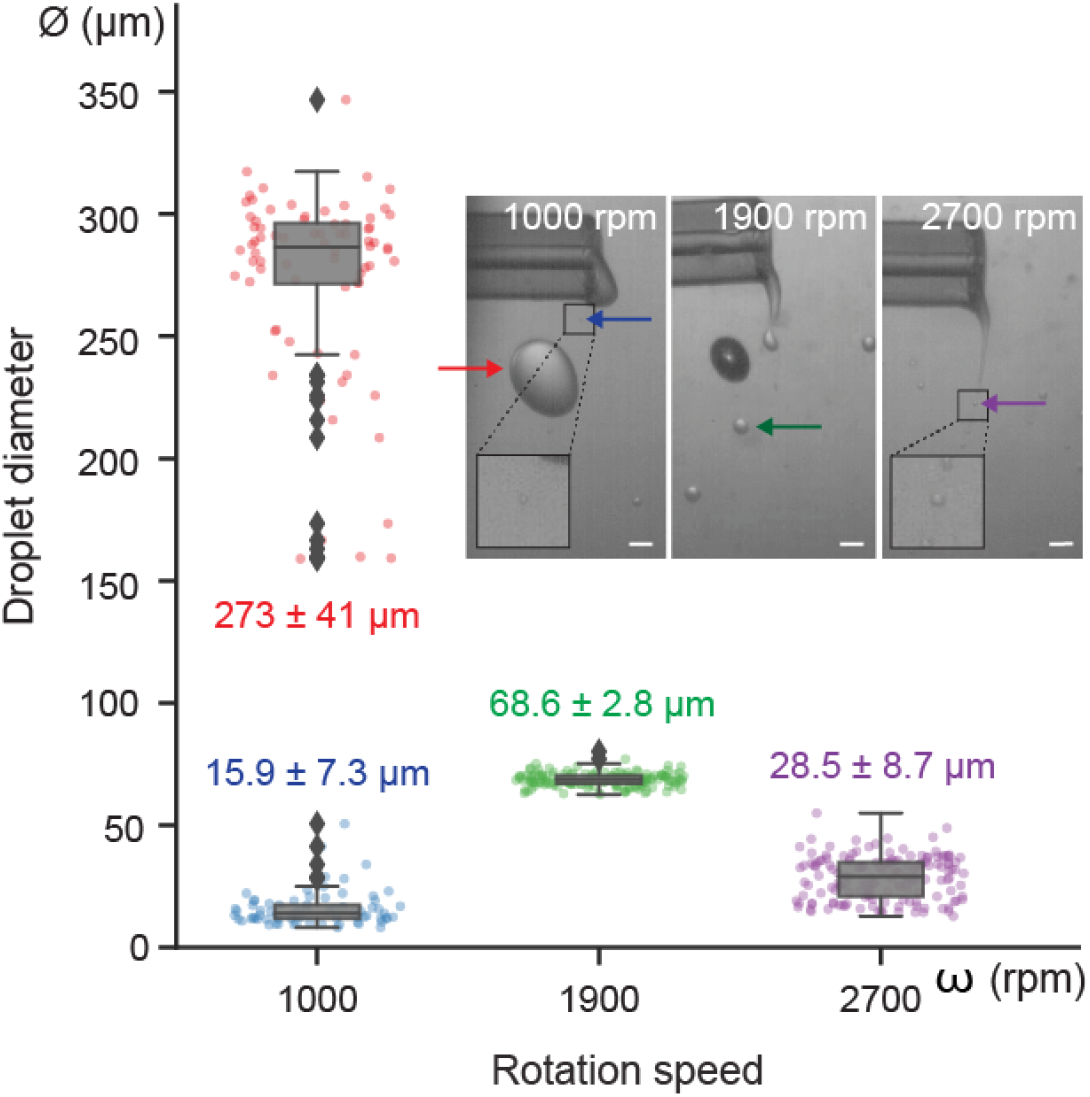
Size distributions of droplets for different rotation speeds. (single column) Boxplots of droplet diameter ϕ at rotation speeds ω of 1000 rpm, 1900 rpm, and 2700 rpm (n = 148, 152, and 157, respectively, for N=1). Individual data points indicate single droplets and boxplots indicate medians and quartiles, while outliers are marked with individual diamond shapes. A rotation speed of 1000 rpm resulted in two distinct droplet populations: large droplets of mean diameter ϕ (273 ± 41) μm (red) and satellite droplets of mean diameter ϕ (15.9 ± 7.3) μm (blue). A rotation speed of 1900 rpm resulted in the narrowest distribution, with a mean droplet diameter ϕ of (68.6 ± 2.8) μm (green). 2700 rpm resulted in the smallest droplet sizes, with a mean diameter ϕ of (28.5 ± 8.7) μm (purple). Inset: representative field-of-views for the different rotation speeds indicating the formed droplets with arrows. Scale bars indicate 100 µm.

While a rotation speed ω of 1900 rpm resulted in the narrowest droplet size distribution of all explored rotation speeds, interestingly, a rotation speed of 1000 rpm resulted in two distinct populations (Figure 3): one primary population of droplets with a mean diameter ϕ of (273 ± 41) μm and a secondary population consisting of smaller droplets with a mean diameter ϕ of (15.9 ± 7.3) μm. Occasionally, this periodic droplet formation of large and small droplets was disrupted when e.g. a droplet merged with the liquid stream or collided with the capillary. Inspecting the videos more closely, we found that the observed population of small droplets are satellite droplets, produced when a bigger droplet breaks off from the main liquid thread at the tip of the capillary (SI Mov. 3). Such satellite droplets have previously been observed in many breakup configurations, from T-junctions to the breakup of droplets in pure shear.^39^ While we did not observe any satellite formation for rotation speeds > 1000 rpm, this may be due to our limited optical and temporal resolution: the satellite droplets observed for 1000 rpm (diameter ∼15 μm) were at the limits of our image resolution, droplets of any smaller diameter were too small to be identified and measured with sufficient certainty (see method section for further details). It is therefore possible that satellite droplets of all sizes, within the size range of the final GUVs (1-20 μm), are also formed, but not detected by our imaging setup.

In addition to the small satellite droplets we observed at 1000 rpm, smaller droplets could theoretically also be formed when larger droplets break up due to shear forces generated in the flow by the rapid relative motion of the bottom wall of the rotational chamber with respect to the capillary. Droplets formed at the tip of the capillary are entrained by the flow in the rotation direction at a high velocity of *U* ≈ Ω*R*_*i*_ ≈ 1*ms*^− 1^ compared to the slow radial motion *Ur* = ρ_*i*_ − ρo)*a*^2^Ω^2^*R*_*i*_ /*µ* ≈ 3 *cms*^− 1^, determined by the balance between centrifugal and viscous forces. These droplets therefore interact with the wake left behind the capillary for several rotations. In the wake, the characteristic shear rate 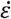 scales with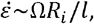, where the characteristic length scale *l* for shear around the capillary will range between the outer diameter of the glass capillary ≈ 0.5*mm* and the distance between the capillary and the bottom of the flow chamber ≈ 5*mm*. One can define another capillary number as 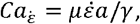, where a is the radius of the droplet.^40,41^ This number characterizes the relative magnitude of the viscous shear forces due to the shear rate c and the surface tension forces. 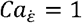 corresponds to a condition where the smallest droplets cannot be further broken up by the shear^40,41^ and yields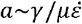 . Knowing *h*∼ 0.5 *cm*, we find that the interfacial tension of the monolayer at the inner solution/oil interface needs to be approximately γ∼10^− 5^ − 10^− 6^ *Nm*^− 1^ to produce droplets of *a*∼5 µm, equivalent to the final GUV size. This value for an interfacial tension at an aqueous/oil interface is extremely low and not expected, even in presence of surfactants or lipids. For reference, the interfacial surface tension between two miscible liquids is of the order 10^−6^ *Nm*^−1^_42_. Hence, we conclude that it is unlikely GUV-sized droplets form by shear force-induced droplet breakup after droplet formation at the capillary orifice.

### Protein in the inner solution affects viscosity and lipid adsorption

Next, we set out to study the effect of proteins on droplet formation at the capillary orifice. It is well known that encapsulation of more complex solute mixtures, such as proteins and their associated buffers, leads to a decreased yield and variable encapsulation efficiencies^31,43^. For cDICE specifically, it has been reported that the yield decreased at high protein concentration^44^, yet it is still unknown why this is the case. We chose to investigate the effects of actin and tubulin, due to the widespread efforts for cytoskeletal reconstitution inside GUVs.

Upon addition of either protein, droplet breakup at the capillary orifice also occurred at the tip of the liquid stream exiting the capillary. However, the oscillations of the liquid stream in the wake of the capillary were significantly reduced (Fig. 4a, SI Mov. 4-7). Remarkably, in the case of tubulin, the liquid stream displayed a tendency to adhere to the air-oil interface. To explain these observations, we characterized the inner solution. We looked into both the physical properties, i.e. dynamic viscosity as determined by bulk shear rheology, and physicochemical properties, specifically lipid adsorption rate determined from pendant drop tensiometry.

**Figure 4.**
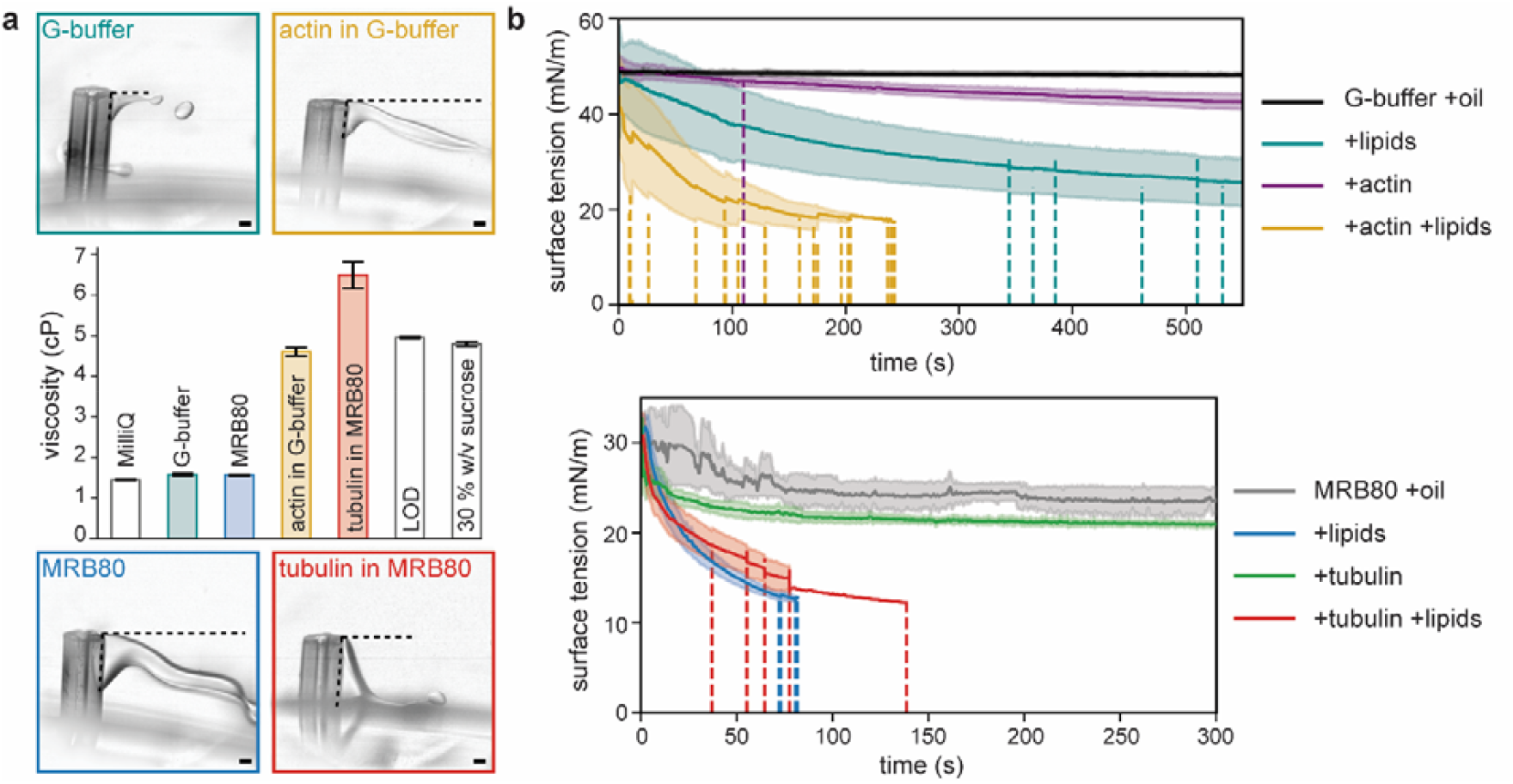
Effect of protein on aqueous solution properties. **a**. Representative field-of-views of droplet formation at the capillary orifice for different protein solutions. Horizontal dotted lines indicate the liquid filament length just before the drop breaks off, while vertical dotted lines along the capillary indicate the extent of external capillary surface wetted by the aqueous solution. Images are background subtracted for better contrast. Scale bars indicate 100 µm. Middle: Dynamic viscosity measured using a parallel plate rheometer for different buffers (G-buffer with 6.5 % v/v OptiPrep™, MRB80 with 1.75 % w/v sucrose) and protein solutions (actin and tubulin), along with water (MilliQ), lipid-in-oil dispersion (LOD) and 30 % w/v sucrose solution in MRB80 for reference. Error bars represent standard deviation. **b**. Interfacial tension kinetics measured using pendant-drop tensiometry for different combinations of aqueous and oil solutions; G-buffer and actin (top), and MRB80 and tubulin (bottom). Solid lines represent the average values and the shaded region corresponds to standard deviation. The vertical dotted lines represent the event of falling of a drop and truncation of data.

In presence of actin and tubulin, the dynamic viscosity increased with respect to its accompanying buffer, G-buffer and MRB80 buffer respectively (Fig. 4a). For actin (1 μM in G-buffer, 6.5 % v/v OptiPrep™), an almost threefold increase from 1.58 cP to 4.61 cP was observed (Fig. 4a, yellow bar), while for tubulin (33.33 μM in MRB80 buffer, 1.75 % w/v sucrose), the viscosity increased fourfold from 1.57 cP to 6.49 cP (Fig. 4a, red bar). All solutions still exhibited Newtonian fluid behaviour. Important to note is that the used concentrations of added proteins remained within the micromolar range and are widely used in the field. Interestingly, the viscosity of the inner solution containing protein was similar to the viscosity of the continuous phase, i.e. the surrounding lipid-in-oil dispersion (Fig. 4a, middle ‘LOD’ bar). The fragmentation of the liquid filament into droplets at the end of the capillary is the consequence of complex instabilities beyond the scope of this study. These mechanisms are significantly affected by the viscosity of the inner solution and the increased viscosity due to the added protein will dampen the flow dynamics in the liquid filament. This dissipation in the liquid stream can explain the decrease in the fluctuations observed in the liquid filament exiting the capillary (Fig. 4a, SI Mov. 4-7). Moreover, previous studies on capillary breakup have reported the viscosity to affect the fragmentation pattern and the size distribution of satellite droplets significantly. In particular, the viscosity increase in a liquid filament has been associated with fewer and larger satellite droplets^45,46^. Therefore, proteins included in the inner solution can have a significant impact in the size distribution of droplet formed at the capillary exit. Altogether, these results show a nuanced interplay between the physical properties of the encapsulation solution, varying with its composition even at low protein concentrations, and the fluid dynamic processes that govern droplet breakup.

To investigate how the addition of protein to the inner solution alters the physicochemical properties of the interface, we used pendant drop tensiometry^47^ to study lipid monolayer formation in a controlled environment. We analyzed the lipid adsorption kinetics and interfacial tension dynamics of the water-oil interface for different encapsulation solutions, mimicking droplet formation at the capillary orifice. It has been shown that proteins spontaneously adsorb at the oil-water interface and their behaviour cannot unequivocally be attributed to a single protein property, with thermodynamic stability, structural properties, and concentration all being contributing factors.^48,49^ Particularly, actin has been shown to exhibit surface activity in a charge-dependent manner, influenced by both lipid and buffer composition, with a more pronounced effect observed for the filamentous form compared to actin monomers.^50–52^

Upon adding 4 μM actin to the inner solution, a pronounced decline in interfacial tension was observed (Fig. 4b, purple curve), with some droplets detaching before the end of the experiment (Fig. 4b, dashed lines). This trend was consistent for tubulin (Fig. 4b, green curve). To examine the roles of actin and tubulin as surface-active agents in the interfacial tension, we then compared the interfacial tension dynamics against a lipid-free oil dispersion. Both actin and tubulin had only a marginal impact on interfacial tension when compared to the protein-free condition (Fig. 4b, black curve vs purple curve and grey curve vs green curve). Interestingly, while actin and lipids individually at the interface exhibited slow kinetics, their combined presence displayed an accelerated decrease (Fig.4b, yellow curve), suggesting a synergistic effect beyond mere additivity. We found this effect could not be countered via electrostatic or steric repulsion (*i*.*e*. presence of charged or PEGylated lipids, respectively, SI Fig. 3). These results imply that actin could, for example, quickly cover the surface of the droplets traversing the lipid-in-oil dispersion, potentially impeding lipid monolayer formation and/or monolayer zipping. However, the full extent of this synergistic effect is yet to be uncovered. Furthermore, these results underscore the importance of the compositions of both inner solution and lipid-in-oil dispersion, as both affect mono- and bilayer formation.

### GUV formation at the oil-water interface seems size selective

Droplet formation in cDICE occurs on extremely short timescales; for the default conditions (i.e. 1900 rpm, 25 uLmin^−1^), we observed droplets of approximately ∼70 µm in diameter being sheared off at a frequency of ∼2500 Hz. Theoretically, given a total encapsulation volume of 100 µL, > 500,000 droplets are formed during a single experiment. Interestingly, this number does not correspond to the final number of GUVs produced using cDICE, as reported in other publications (∼ 1000 GUVs^31^).

Furthermore, if those droplets larger than the finally observed GUVs (i.e. non satellite droplets, ∼70 µm for the default conditions) do not subsequently shear to form smaller droplets as discussed above, these two observations together indicate a sub-optimal GUV formation process downstream, whereby most droplets do not convert into GUVs at the oil-water interface and potential additional hidden mechanisms generating smaller droplets.

To look more closely at droplet-to-GUV conversion into GUVs in cDICE, we imaged the oil-water interface where the final step of GUV formation in cDICE occurs: droplets transfer through the oil-water interface and two monolayers fuse together to form a bilayer (Fig. 5a). As postulated by Abkarian *et al*.^30^, the two monolayers can also form a pore, thereby causing the droplet to burst, resulting in no GUV being formed. We note that when we collected GUVs in cDICE experiments, we observed that the outer solution after GUV generation also contained components of the inner solution, in agreement with the suggestion that a fraction of droplets burst at the oil-water interface.

**Figure 5.**
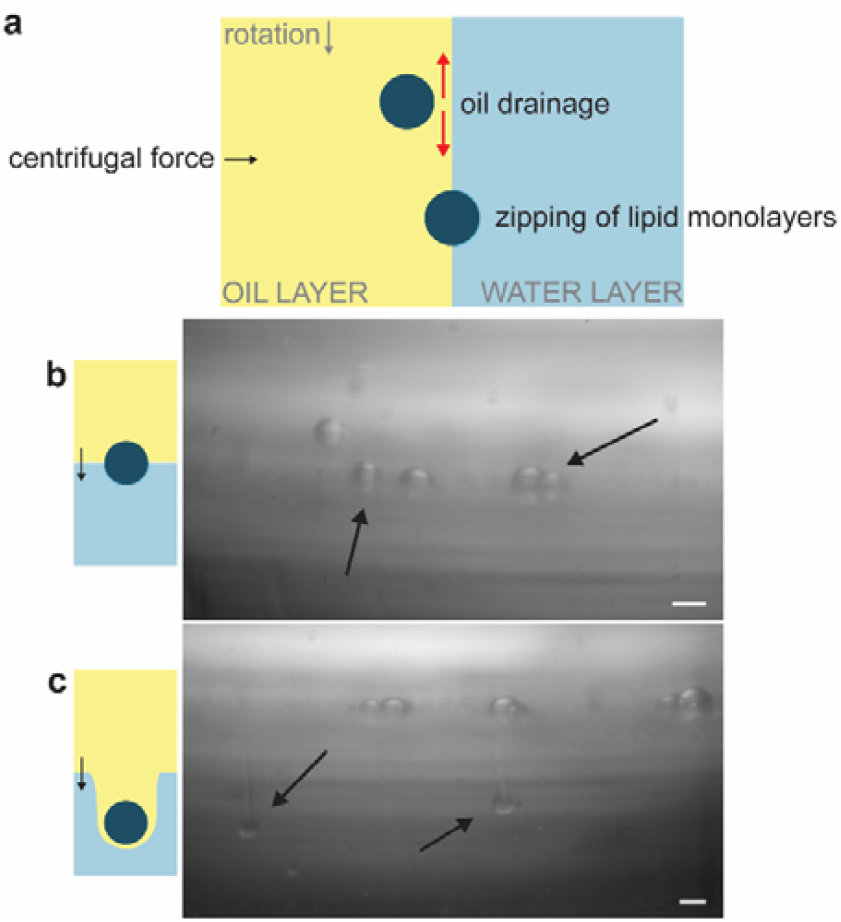
Droplet transfer through the oil-water interface is suboptimal. (single column) **a**. The lipid-in-oil dispersion in between the approaching lipid monolayer-covered droplets and the oil-water interface needs to be drained for the two monolayers to zip together and successful GUV formation to occur. **b**. When droplets are not fully covered by a lipid monolayer when reaching the oil-water interface, successful transfer cannot occur and instead, stationary, semi-transferred droplets are observed. Scale bar indicates 100 μm. **c**. When drainage of the oil layer between the approaching droplet and oil-water interface is insufficient, the formation of comet tails can be observed: a droplet distorts the oil-water interface and drags the lipid-in-oil dispersion into the outer aqueous solution, hindering successful GUV formation. The Scale bar indicates 100 μm.

In our experiments, we unfortunately did not observe a clear transfer of droplets through the interface nor bursting of droplets, possibly because resolving GUV-sized droplets at the interface was not feasible with the limited imaging contrast of standard bright field illumination. Instead, we made two other, striking observations. First, we observed droplets several orders of magnitude larger than the typical size of GUVs which were stationary on the oil-water interface (Fig. 5b, SI Mov. 8). These stationary droplets showed a decreased contrast on the side of the outer aqueous phase, suggesting partial transfer across the interface. Since the transfer time of a droplet to the oil-water interface is inversely proportional to the radius of the droplets squared^30^, it is possible that the flight time of these larger droplets is too short for lipids to fully adsorb on the interface. Consequently, no zipping mechanism is possible, leading to these larger droplets crowding the interface, as we observed in our video recordings.

A second observation was the formation of comet tails (Fig. 5c): droplets that did pass the interface dragged a tail of the oil solution into the outer aqueous solution, likely because the oil did not drain quickly enough and thus prevented monolayer fusion. Due to the difference in contrast with the outer aqueous phase, we infer that oil is still present between the part of the interface dragged into the outer phase and the droplet. The Bond number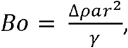, where *a* is the acceleration and *r* the radius of droplet, represents the ratio of centrifugal force to surface tension force. For these large droplets, *Bo* is on the order of 1, meaning they will deform the interface, as observed in our video recordings, and drag the oil phase into the outer aqueous phase. This results in the observed comet tail formation and no GUV formation from the droplets undergoing this process.

As we find that the addition of protein to our inner solution significantly alters the characteristics of the solution and affects droplet formation at the capillary orifice, we asked how the increased viscosity and altered lipid adsorption kinetics might impact the transfer of droplets through the oil-water interface. The accelerated lipid adsorption due to the addition of protein does not lead to a decreased flight time of the droplets, thereby not adversely affecting droplet transfer or monolayer zipping. On the other hand, the increased viscosity of the inner solution could influence the timescale of the drainage of the lubrication film, i.e. the lipid-in-oil dispersion in between the droplet and the oil-water interface, required for successful monolayer zipping. Furthermore, the increased viscosity could reduce the flow caused by Marangoni stresses, which play a role in facilitating the zipping process.^30^

An approximate breakthrough condition for spherical objects of radius a to pass through an interface of interfacial tension γ is (ρ_*i*_ − ρ*o*)Ω^2^*Roa*^2^/γ ≥ 3/2, where *Ro* is the distance between the axis of rotation and the location of the oil-outer solution interface.^53^ For small droplets of radius *a* ∼ 5µ*m* to cross the interface, a low surface tension on the order of γ ∼10^− 6^*Nm*^− 1^ is required. Such low surface tension has been reported for lipid bilayers^54^ and therefore if such small droplets are present in the oil phase, they can cross the interface to form GUVs. It should be emphasized that the breakthrough condition sets a criterion for the smallest droplet that can cross the interface. Any droplet larger than 10 *µ*m in diameter would be expected to cross the interface as well and form larger GUVs. The fact that we do not observe GUVs of larger diameters than ∼20 μm^31^, but do observe large droplets at the oil-water interface, suggests that the upper size limit for GUV formation might be controlled by membrane zipping and/or lipid coverage of the droplet/interface. Insufficient lipid coverage could, for example, lead to droplet/GUV shrinkage during GUV forming until the lipid density to form a bilayer is reached, thereby resulting in smaller GUVs than originally produced droplets.

Comparing cDICE GUV size distributions to those obtained by eDICE, a recent adaptation of cDICE where the droplets are generated by vortexing, pipetting, or scraping, instead of a capillary, but transferred through a second interface in a rotational chamber, identical to cDICE. Interestingly, we noticed that the final GUV size distributions were similar for the two methods^55^, despite vastly different droplet size distributions were used as a starting point(SI Fig. 4). Furthermore, we found GUV sizes to be remarkably similar for different membrane compositions in eDICE (SI Fig. 5). Taken together, these cDICE and eDICE results indicate a yet unknown mechanism for size-selectiveness at the oil-water interface that promotes the production of similarly-sized GUVs for a wide distribution of droplet sizes. For example, it is possible that GUVs form at the oil-water interface in cDICE and eDICE by pinching off from larger droplets sitting at the interface. While we did not observe any event like this, we would expect this process to happen on a length scale (and possible time scale) beyond the resolution of our imaging setup.

## Conclusion

In summary, by designing and building a custom imaging setup to visualize droplet formation and droplet interface transfer in cDICE in real-time, we were able to, for the first time, collect direct *in situ* imaging data to further understand the underlying mechanisms governing GUV formation in this technique. We found that droplet formation at the capillary orifice produced droplets that are much larger than the size of the final GUVs. For a capillary diameter of 100 μm, the formation of droplets in cDICE bears some similarities with the formation of droplets at T-junctions in microfluidics, a well-studied phenomenon.^56,57^ In such microfluidic channels, the geometric confinement provided by the channels leads to flow restrictions on the continuous phase at the origin of a squeezing pressure. This pressure promotes droplet break up at much smaller values of *Ca* as compared to our experiments. However, there are similarities in the droplet formation regimes. For example, a decrease in droplet volume for increasing values of *Ca* has been widely reported.^56,57^ These studies have also reported a transition from a breakup droplet formation mechanism for low values of *Ca* to a dripping mechanism at higher *Ca*, whereby a long liquid filament of the dispersed phase forms and droplets pinch off at the end of the filament. This is in contrast to the use of smaller capillary openings in the original cDICE implementation, in the range of 2-25 μm^30^, where the smaller inner diameter of the capillary leads to smaller droplet sizes by a combination of smaller total interfacial force resisting the breakup of the droplet and a smaller Reynolds number. Only as a side process, smaller satellite droplets are being formed. Even smaller droplets, including GUV-sized droplets, could not be resolved due to the limited resolution of our setup. Furthermore, we showed that the addition of protein to the inner solution increases its viscosity and changes interfacial tension dynamics, both directly impacting droplet formation and interface transfer. Imaging of the oil-water interface revealed that droplet transfer is frequently stalled, sub-optimal for large droplets, and exhibits a size-selectivity. This size-selectivity of droplet transfer to GUVs was further confirmed using eDICE, a variant of cDICE where no capillary is used, which yielded similar size distribution despite vastly different droplets as input. Further studies are needed to further elucidate the effect of lipid composition, including cholesterol or charged lipids, and different proteins or protein mixes. We believe the presented results can be of interest not only for cDICE but to other emulsion-based GUV formation methods as well, ultimately contributing to the development of more reliable and efficient methods for GUV production. Our study furthermore emphasizes the need for inter-disciplinary collaboration to fully grasp the intricacies of the processes involved in emulsion-based GUV production methods. Altogether, this research is just a prologue to a larger narrative and we hope it will serve as a stepping stone for future research, enhancing emulsion-based GUV formation along the way.

## Methods

### Design and fabrication of the spinning device

The cDICE device was identical to Van de Cauter *et al*.^31^. An additional opening underneath the spinning chamber was created by removing part of the motor housing. This way, the light source could be placed directly below the spinning chamber to achieve transillumination. The design for the adjusted cDICE device is available on GitHub (https://github.com/GanzingerLab/cDICE_microscope).

### Fabrication of spinning chambers

Transparent, cylindrical chambers, 35 mm in diameter and 10 mm in height, were made from two lids of Petri dishes (Falcon® REF 351008). To create a waterproof, closed chamber, the sides of the two lids were first sanded using sandpaper to create a rough surface after which they were glued together using a thin layer of optical glue (Norland Optical Adhesive 81). After curing of the glue using UV light, the side of the chamber was wrapped with a strip of Parafilm®. The chambers include a circular opening of 15 mm in diameter in the top to allow facile access to the solutions with the capillary.

### General cDICE experimental workflow

While it is possible, and needed, to tweak various operational parameters to encapsulate a particular (non-)biological system in cDICE, we chose to use the parameters established in a recent optimization study by Van de Cauter et al.^31^ as default conditions for cDICE. Specifically, we used a 100 µm diameter capillary, a rotation speed of 1900 rpm, and a flow rate through the capillary of 25 µLmin^-1^. For the lipid-in-oil dispersion, 18:1 1,2-dioleoyl-sn-glycero-3-phophocholine (DOPC) lipids were dispersed using chloroform in a 4:1 ratio silicon oil:mineral oil (silicon oil – viscosity 5 cst (25°C), Sigma-Aldrich; mineral oil – BioReagent, Sigma-Aldrich). A fused silica capillary tubing with polyimide coating (TSP-100375, Molex LLC) was used to inject inner aqueous solutions. The general cDICE experimental workflow and preparation of lipid-in-oil dispersion was based on Van de *Cauter et al*.^31^. The following parameters differed: The volume of the outer solution was increased to 1.07 mL to account for the difference in dimensions between the 3D printed spinning chambers, as used in Van de Cauter et al.^31^, and the Petri dish spinning chambers that were used for imaging experiments, as mentioned above. Room humidity was not controlled during imaging experiments and the chambers were spun for the entirety of the imaging experiments instead of a predetermined time. G-buffer (5 mM tris(hydroxymethyl)aminomethane hydrochloride (Tris-HCl) pH 7.8 and 0.1 mM calcium chloride (CaCl_2_), 0.02 mM adenosine triphosphate (ATP) and 4 mM dithiothreitol (DTT)) with 18.5 % v/v OptiPrep™ was encapsulated in every experiment (to achieve a density difference between the inner and outer aqueous solution), unless specified otherwise. For experiments with silanized capillaries, the tip of the capillary was submerged for one minute in dichlorodimethylsilance (DMDCS) (40140, Sigma Aldrich), before removing excess with nitrogen gas.

### Home-built imaging setup

The light of a single LED (Luxeon V2, 315lm@700mA; used without lens) or a Lumencor light engine (SOLA 6-LCR-SB) was collected by a 200 mm focal length achromatic lens (Thorlabs AC254-200-A-ML; lens mount: Thorlabs CXY1). The setup was equipped with a 4X or 10X objective (Nikon Plan Fluor 4x/0.13 PhL DL and Nikon Plan Fluor 10x/0.30 ∞/0.17 WD 16, respectively) that was mounted on a Z-stage (Thorlabs CT1; adapter: Thorlabs SM1A10). X/Y motion control was provided by two translational stages with a step size of 25 mm (Thorlabs PT1). Images were recorded using a high-speed camera (Kron Technologies Chronos 2.1-HD and Photron FASTCAM SA4) which was mounted on the setup using a custom-designed 3D printed construction. The full setup was mounted on a Thorlabs cage system that was mounted on a breadboard (Thorlabs MB1030/M) to easily move the full setup over the cDICE device. The full component list and design plans, including an interactive 3D model of the setup, can be found on GitHub (https://github.com/GanzingerLab/cDICE_microscope).

### Droplet size analysis

Droplet size analysis was performed manually using the Fiji software^58^. The image pixel size was derived from three independent measurements of the capillary opening, accounting for capillary size uncertainty. Triplicate measurements were performed for a subset of each dataset to quantify the measurement error. For each droplet, we then measured both area and diameter, yielding two independent measurements of the droplet diameter with associated error calculated through error propagation. The large pixel size ((2.431 ± 0.105) μm) in comparison to the droplet size characterized, in combination with a measurement error of 2 μm, calculated from measuring a subset of data in triplicate, posed a limit on our analysis of smaller droplets. Additionally, the high speed of the process, resulting in motion blur and droplets quickly moving out of focus, as well as the limited contrast, caused by the small difference in refractive index between the droplets and the surrounding medium (1.333 for water vs 1.403 for silicone oil), makes it difficult to distinguish the droplets from the background in the video recordings. Data visualization was achieved through Python-generated plots. The frequency was estimated using the mean droplet size and the flow rate of the inner solution. Note that for the analysis of droplet size and frequency, we used video recordings in which the fluid tail did not adhere to the capillary surface (one experiment per condition).

### Viscosity measurements

The dynamic viscosities of the solutions were measured on a Kinexus Malvern Pro rheometer. A stainless steel plate-plate geometry with 40 and 20 mm radius were used for buffer solutions and protein-containing solutions respectively. Viscosity was measured every 5 seconds as a function of shear rate with a 2 min logarithmic viscometry ramp from 0.5 s^− 1^ to 100 s^− 1^. As expected for simple viscous liquid, viscosities for higher shear rate were constant. The values at 100 s^− 1^ were used to calculate the reported viscosity of each solution. MRB80 buffers consists of 80 mM piperazine-N,NC-bis(2-ethanesulfonic acid) (PIPES) pH 6.8, 4 mM magnesium chloride (MgCl_2_), and 1 mM ethylene glycol-bis(β-aminoethyl ether)-N,N,N⍰,N⍰-tetraacetic acid (EGTA).

### Tensiometry measurements

The pendant drop measurements were performed using a DSA 30S drop shape analyser (Kruss, Germany) and analysed with the Kruss Advanced software. Experimental conditions for G-buffer and actin-containing solutions were as described in Van de Cauter *et al*.^31^, while changes for MRB80 buffer and tubulin-containing solutions are described below. Initially a 2 μL droplet of aqueous solution is drawn in a lipid-in-oil dispersion containing glass cuvette (Hellma Analytics) and then volume of the droplet is adjusted to 8 μL using an automated dosing system from a hanging glass syringe with needle diameter of 0.313 mm (Hamilton). As soon as the droplet reached its final volume, droplet was analysed (for 300 seconds at 25 fps for solutions containing tubulin and lipids and at 5 fps for rest of the solutions) by automatic contour detection and fitted with the Young-Laplace equation to yield the interfacial tension. The densities of lipid oil solution (0.8685 mg/ml), G-buffer with 18.5 %v/v OptiPrep™ (1.0574 mg/ml), and MRB80 with 1.75 %w/v Sucrose (1.0066 mg/ml) were used in the interfacial tension calculations. These densities were measured by weighing 1 ml of solution. For G-buffer with OptiPrep™, the density was estimated using the volume-weighted-mean. The surface tension values were smoothened with a rolling-mean of 1 sec. Room humidity was not controlled. In several experiments, interfacial tension decreased very rapidly (abnormally), causing the droplet to detach as soon as they are formed. These measurements were discarded from analysis.

## Supporting information

SI Mov 1

SI Mov 2

SI Mov 3

SI Mov 4

SI Mov 5

SI Mov 6

SI Mov 7

SI Mov 8

## Acknowledgements

We thank Roy Hoitink for his contributions during the initial exploration of the imaging setup, SaFyre Reese for experimental help during parameter screening, and Irene Istúriz Petitjean for help with rheology experiments. We would also like to thank Dr. Arjen Jakobi for generously lending us the Chronos 2.1-HD camera, and Iliya Cerjak and Bob Krijger for technical help with the design of the imaging setup and the light source, respectively. We acknowledge financial support by the “BaSyC – Building a Synthetic Cell” Gravitation grant (024.003.019) of The Netherlands Ministry of Education, Culture and Science (OCW) and The Netherlands Organization for Scientific Research (NWO) (M.D. and G.H.K.) and NWO-WISE funding (K.A.G.). Part of this research was funded by a Pieter Langerhausen Stipendium of the Koninklijke Hollandsche Maatschappij der Wetenschappen (K.A.G).

