## Supplementary material for "High-speed imaging of giant unilamellar vesicle formation in cDICE": SI

**Document containing:**

- **SI Figure 1**
- **SI Figure 2**
- **SI Figure 3**
- **SI Figure 4**
- **SI Figure 5**
- **SI Movie 1**
- **SI Movie 2**
- **SI Movie 3**
- **SI Movie 4**
- **SI Movie 5**
- **SI Movie 6**
- **SI Movie 7**
- **SI Movie 8**

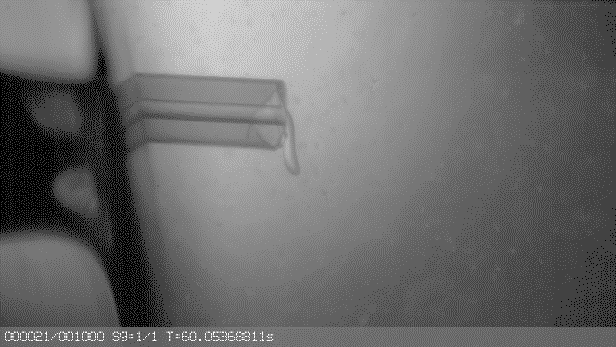

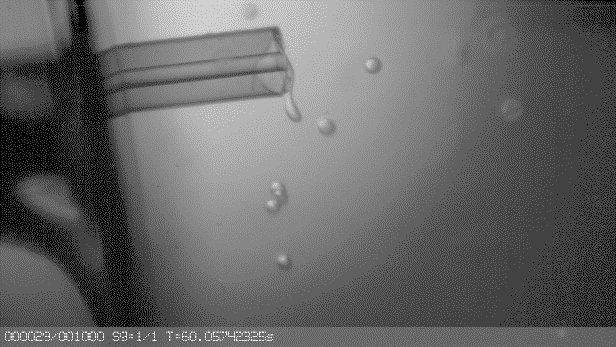

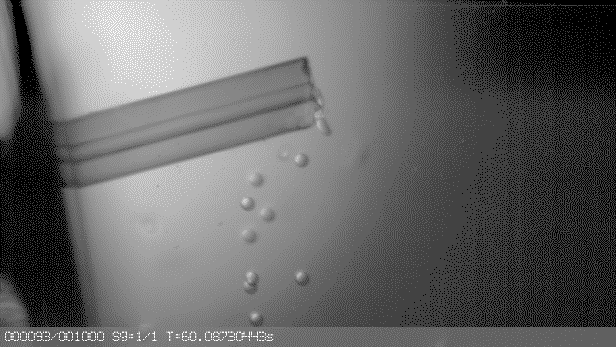

**SI Figure 1. Silanization of the capillary prevented attachment of the liquid thread to the capillary.**

Representative field-of-views, differences can be seen in insertion angle, insertion depth, and capillary orifice (i.e. cut finish and coating chipping off). In all three cases, the capillary was silanized, and the liquid stream did not adhere to the capillary. The capillary was silanized using dichlorodimethylsilane (40140, Sigma Aldrich) by submerging the tip of the capillary for one minute, before removing excess with nitrogen gas. The capillary opening is 100 µm.

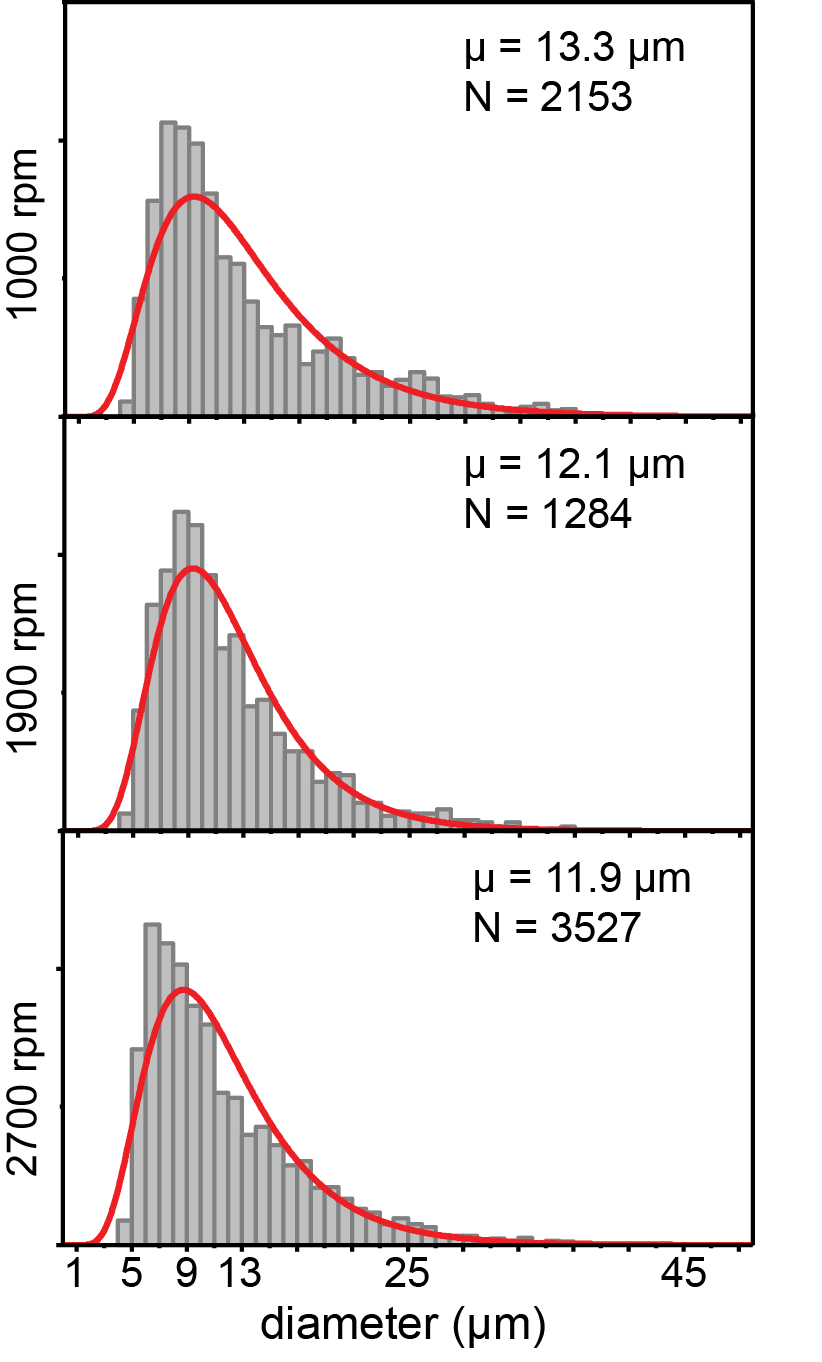

**SI Figure 2. Size distributions of GUVs for different rotation speeds.**

Size distribution of GUVs made at rotation speeds ω of 1000 rpm, 1900 rpm, and 2700 rpm. The individual graphs represent pooled data for three experiments. The distributions are fitted to a log-normal function(red curves). Reprinted (adapted) with permission from Van de Cauter et al 2021.

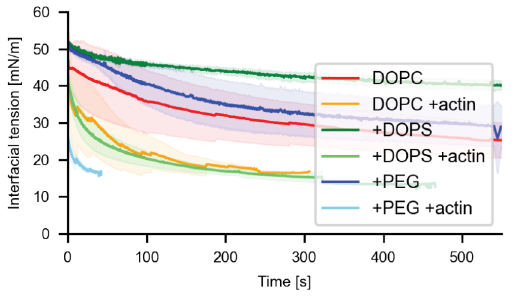

**SI Figure 3. Interfacial tension evolution with and without actin using different lipid compositions.**

Each curve represents the average over N measurements with the shaded region being the standard deviation. Only DOPC against inner aqueous solution (G-buffer with 18.5 % v/v OptiPrep™) without (red line, N=21) and with (orange line, N=19) 4.4 µM actin, DOPC with 20% DOPS against inner aqueous solution without (dark green line, N=2) and with (light green line, N=3) 4.4 µM actin, and finally DOPC with 5% PEG 2000 DOPE measured against inner aqueous solution without (dark blue line, N=3) and with 4.4 μM actin (light blue line, N=3). Total lipid concentration is 0.2 mg/mL.

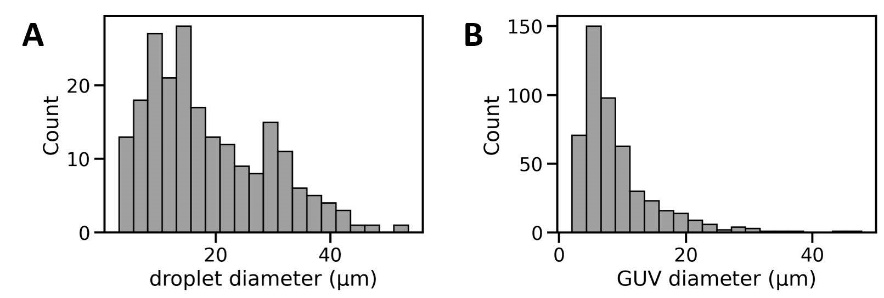

**SI Figure 4. Size distribution of droplets and GUVs in eDICE.**

**a.** Histogram of the diameters of droplets generated in the first step of eDICE (N = 213 droplets from 2 separate experiments). Droplets contained 4.4 µM actin in F-buffer with 6.5 v/v% OptiPrep™ and were stabilized by surfactants in addition to lipids. **b.** Histogram of GUV sizes generated by eDICE (N = 494 GUVs from 4 separate experiments).

eDICE GUVs were prepared as described in Baldauf *et al.*^1^ Emulsion droplets were produced in an identical fashion to those produced in the intermediate GUV preparation step in eDICE, where we emulsified disperse phase into 1 mL of oil phase by mechanical agitation. To allow us to image the aqueous droplets in oil, we additionally dissolved 2 % v/v Span80 (Sigma Aldrich) in the oil phase to stabilize the interface^2^. This addition of surfactant was necessary to keep the droplets from coalescing before or during the ~ 30 min necessary for imaging, but it changes the interfacial properties and may thus have an impact on the droplet size distribution we generate. The addition of extra surfactant has been shown to decrease the average size of emulsion droplets generated in turbulent flow^3^, so our measurements likely underestimate the true size of droplets generated during the eDICE GUV formation process. Epifluorescence images of water-in-oil droplets and GUVs were acquired on an inverted Nikon Ti Eclipse microscope equipped with a 60x water immersion objective (CFI Plan Apochromat VC), a digital CMOS camera (Orca Flash 4.0), and an LED light source (Lumencor Spectra Pad X). Phase contrast images were acquired on the same Nikon Ti microscope.

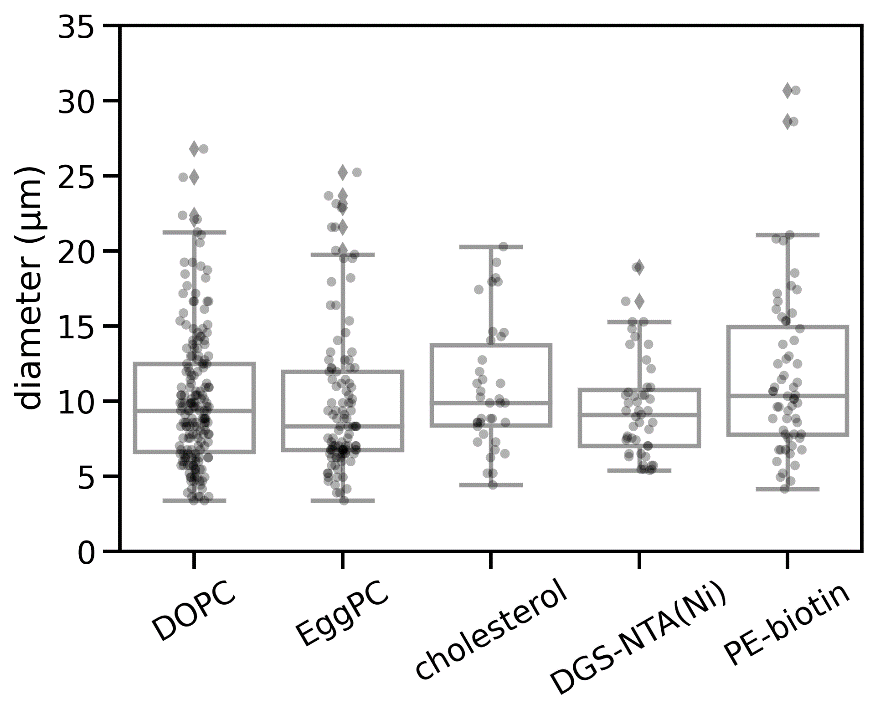

| **Composition** | **Mean(d)** | **Std(d)** | **n** | **Imaged by** |
| --- | --- | --- | --- | --- |
| 100 % DOPC | 10.08 | 4.49 | 187 | Phase contrast |
| 100 % EggPC | 9.844 | 4.89 | 108 | Phase contrast |
| 80:20 DOPC:cholesterol | 10.88 | 4.20 | 38 | Phase contrast |
| 95:5 DOPC:DGS-NTA(Ni) | 9.522 | 3.32 | 47 | Widefield |
| 95:5 DOPC:bioPE | 11.68 | 5.43 | 60 | Widefield |

**SI Figure 5. Size distribution for GUVs produced using eDICE with various membrane composition.**

GUVs with different membrane composition produced by eDICE (see table above; GUVs were prepared as described in Baldauf *et al.*^1^). The membranes for widefield imaging also contained 0.05 % Cy5-labeled lipids. GUVs with 5 % Biotin-PE in the membrane also encapsulated 0.88 µM Atto 488-conjugated streptavidin (Sigma Aldrich), all others are without encapsulated proteins. Sizes were measured manually in Fiji, fitting circles to max projections of a Z-stack. Box plots comparing the GUV sizes are displayed on the graph, the distributions are not statistically different (p=0.070 by one-way ANOVA). Epifluorescence images of GUVs were acquired on an inverted Nikon Ti Eclipse microscope equipped with a 60x water immersion objective (CFI Plan Apochromat VC), a digital CMOS camera (Orca Flash 4.0), and an LED light source (Lumencor Spectra Pad X). Phase contrast images were acquired on the same Nikon Ti microscope.

**SI Movie 1. Droplet formation at the capillary orifice at a rotation speed of 1900 rpm.**

Video recordings of droplet formation at the 100 µm diameter fused silica capillary orifice at a rotation speed of 1900 rpm. The playback speed is 10 fps for a total of 250 frames.

**SI Movie 2. Droplet formation at the capillary orifice at a rotation speed of 2700 rpm.**

Video recordings of droplet formation at the 100 µm diameter fused silica capillary orifice at a rotation speed of 2700 rpm. The playback speed is 10 fps for a total of 250 frames.

**SI Movie 3. Droplet formation at the capillary orifice at a rotation speed of 1000 rpm.**

Video recordings of droplet formation at the 100 µm diameter fused silica capillary orifice at a rotation speed of 1900 rpm. The playback speed is 10 fps for a total of 250 frames. The video has 3.5x zoom inset from region under the capillary for better visualization of the satellite droplets.

**SI Movie 4. Droplet formation at the capillary orifice for G-buffer.**

Video recordings of droplet formation at the 100 µm diameter fused silica capillary orifice with G-buffer as inner solution. The playback speed is 10 fps.

**SI Movie 5. Droplet formation at the capillary orifice for actin in G-buffer.**

Video recordings of droplet formation at the 100 µm diameter fused silica capillary orifice with actin in G-buffer as inner solution. The playback speed is 10 fps.

**SI Movie 6. Droplet formation at the capillary orifice for MRB80 buffer.**

Video recordings of droplet formation at the 100 µm diameter fused silica capillary orifice with MRB80 buffer as inner solution. The playback speed is 10 fps.

**SI Movie 7. Droplet formation at the capillary orifice for tubulin in MRB80 buffer.**

Video recordings of droplet formation at the 100 µm diameter fused silica capillary orifice with tubulin in MRB80 buffer as inner solution. The playback speed is 10 fps.

**SI Movie 8. Oil-water interface**

Video recording of the oil-water interface. The playback speed is 2 fps for a total of 250 frames.
